# Rapid Sequential *in Situ* Multiplexing With DNA-Exchange-Imaging

**DOI:** 10.1101/112227

**Authors:** Yu Wang, Johannes B. Woehrstein, Noah Donoghue, Mingjie Dai, Maier S. Avendaño, Ron C.J. Schackmann, Jason J. Zoeller, Shan Shan H. Wang, Paul W. Tillberg, Demian Park, Sylvain W. Lapan, Edward S. Boyden, Joan S. Brugge, Pascal S. Kaeser, George M. Church, Sarit S. Agasti, Ralf Jungmann, Peng Yin

## Abstract

To decipher the molecular mechanism of biological function, it is critical to map the molecular composition of individual cells in the context of their biological environment *in situ*. Immunofluorescence (IF) provides specific labeling for molecular profiling. However, conventional IF methods have finite multiplexing capabilities due to spectral overlap of the fluorophores. Various sequential imaging methods have been developed to circumvent this spectral limit, but are not widely adopted due to the common limitation of requiring multi-rounds of slow (typically over 2 hours at room temperature to overnight at 4 °C in practice) immunostaining. DNA-Exchange-Imaging is a practical platform for rapid *in situ* spectrally-unlimited multiplexing. This technique overcomes speed restrictions by allowing for single-step immunostaining with DNA-barcoded antibodies, followed by rapid (less than 10 minutes) buffer exchange of fluorophore-bearing DNA imager strands. By eliminating the need for multiple rounds of immunostaining, DEI enables rapid spectrally unlimited sequential imaging. The programmability of DNA-Exchange-Imaging allows us to further adapt it to diverse microscopy platforms (with Exchange-Confocal, Exchange-SIM, Exchange-STED, and Exchange-PAINT demonstrated here), achieving highly multiplexed *in situ* protein visualization in diverse samples (including neuronal and tumor cells as well as fresh-frozen or paraffin-embedded tissue sections) and at multiple desired resolution scales (from ~300 nm down to sub-20-nm). Validation highlights include 8-target imaging using single-channel Exchange-Confocal in tens of micron thick retina tissue sections in 2-3 hours (as compared to days required in principle by previous methods using comparable equipment), and 8-target super-resolution imaging with ~20 nm resolution using Exchange-PAINT in primary neurons. These results collectively suggest DNA-Exchange as a versatile, practical platform for rapid, highly multiplexed *in situ* imaging, potentially enabling new applications ranging from basic science, to drug discovery, and to clinical pathology.

## Introduction

Fluorescence microscopy has become a standard tool to characterize specimens in biological and biomedical studies. One of its advantages is the widespread availability of protein-specific labeling reagents such as antibodies. However, while dye-labeled antibodies allow for easy target labeling, the spectral overlap of multiple fluorophores leads to limited multiplexing capabilities (e.g. typically no more than 4 targets). This shortcoming currently prevents studies targeted towards investigating network-wide changes in single cells and tissues using fluorescence microscopy. Various techniques, including ‘dye-cycling’ by repeated antibody staining^1,2,3,4,5,6,7,8^, multiplexed ion beam imaging (MIBI)^9,10,11^, spectrally resolved stochastic reconstruction microscopy (SR-STORM)^12^, as well as others^13,14,15,16^, have been developed to overcome current limitations for multi-target detection, enabling highly multiplexed imaging studies (see **Supplementary Table 1** for a detailed comparison of these different techniques). However, these techniques have thus far not been broadly adopted due to practical limitations: they are typically time intensive (e.g. due to repeated antibody staining as in current dye-cycling techniques, with each round of staining taking hours at room temperature and preferentially overnight at 4 °C for optimal labeling), and/or difficult to be directly implemented into current widely available microscope systems because specialized instruments are often required (e.g. as MIBI and SR-STORM).

To overcome current limitations, we introduce DNA-Exchange-Imaging (DEI), a generalization of our previously developed Exchange-PAINT^17^ technique, providing a fast and practical method to perform highly multiplexed fluorescence imaging using standard, commercially available microscopy platforms. In DEI, we employ DNA-barcoded antibodies – instead of dye-labeled antibodies – that are conjugated with short DNA oligos (typically 9–10 nucleotides long for implementations in this paper) called docking strands^18^. Multiplexed protein target labeling is performed efficiently by single-step simultaneous immunostaining with antibodies carrying orthogonal DNA docking strands, followed by image acquisition where dye-labeled complementary imager strands are applied sequentially via rapid buffer exchange (**Fig. 1**).

**Figure 1.**
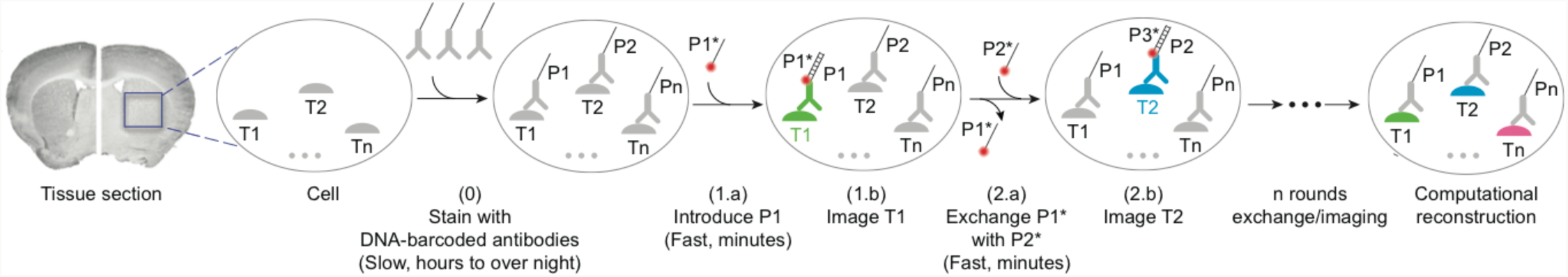
DNA-Exchange-Imaging. Distinct targets (T1, T2, …, Tn) are labeled using corresponding antibodies conjugated to orthogonal DNA docking strands (P1, P2, …, Pn) in a single step. Imager strands (P1*, P2, …, Pn*) are sequentially introduced to visualize target signals. The imager strands are washed away rapidly using low ionic strength buffer after each round of imaging. After imaging, all images are computationally registered and a final merged image is reconstructed by assigning pseudo-colors to each target image.

We have previously demonstrated DEI in the form of Exchange-PAINT^17^. In this paper, we further demonstrate that the DNA-Exchange-Imaging principle can be generalized to other diverse super-resolution microscopy systems, including SIM and STED. Additionally, we extend DEI to standard resolution confocal microscopes (which are widely available in common biological labs) for rapid, spectrally unlimited *in situ* multiplexing in cells and, importantly, in thick tissues (as compared to thin layer of cells in our previous^17^ and recent^19^ super-resolution Exchange-PAINT work). Unlike the fluorescence ‘blinking’ in our previous single molecule-based Exchange-PAINT, the Exchange-Confocal uses pseudo-permanent and dense target labeling with fluorophore-conjugated complementary imager strands, thus permitting rapid image acquisition (typically <1 s exposure time) and deeper sample penetration (tens of micrometers versus a few hundred nanometers in PAINT). In addition to providing a rapid and simple multiplexed imaging method, Exchange-Confocal, as well as other DEI methods, enables easy autofluorescence correction, and is naturally chromatic aberration-free and photobleaching-resistant (**Supplementary Fig.1**).

## Results

### Diffraction-limited Exchange-Confocal imaging in primary neuron culture

Exchange-PAINT^17^ has been previously developed to perform multiplexed single-molecule localization-based SR imaging. Despite its superior resolution, its utility is restricted due to its imaging time and depth tradeoff. It requires recording a time-lapse movie of single molecule blinking events for final SR image reconstruction, which typically takes minutes to even hours for a single reconstructed image. In addition, the high signal-to-noise ratio requirement for PAINT imaging, single-molecule-compatible microscopes (usually Total Internal Reflection Fluorescence [TIRF] microscopes) are necessary, limiting the imaging depth to typically a few hundred nanometers above the coverslip. Moreover, diffraction-limited imaging is often sufficient for experiments that only require single-cell resolution (*e.g.* pathological analysis). In SR PAINT imaging, sparse labeling of targets with transiently binding imager strands is required for singlemolecule localization. In contrast, diffraction-limited Exchange-Confocal imaging shown here aims to capture signals from all the molecules of a certain target in a single image frame, which requires pseudo-permanent and dense target labeling with imager strands. To achieve this, we tuned three parameters: imager/docking strand association time, imager strand concentration and camera exposure time. First, we designed imager/docking strand duplexes with higher binding affinity to attain relatively slow dissociation rate (0.2 s^-1^ for a 10 base-pair duplex on average^18^) by increasing the length of the DNA duplex (**Supplementary Fig. 2**). To minimize the number of unoccupied docking sites, we used a high imager strand concentration (*e.g.* 10 nM as compared to 1 nM in single-molecule PAINT applications) to densely label the docking sites for the corresponding target (**Supplementary Fig. 2**). Furthermore, we used longer camera exposure times (typically 50 to 300 ms for a widefield microscope and 500 ms to 5 s for a spinning disk confocal microscope) to minimize unoccupied docking sites and enhance the signal-to-noise ratio.

As a result, we achieved diffraction-limited Exchange-Confocal images with a quality comparable to that of conventional IF methods. To examine signal specificity of Exchange-Confocal, we compared the Exchange-Confocal images with those from conventional IF methods using fluorophore-conjugated antibodies (**Fig. 2a** and **Supplementary Fig. 3**). We labeled synapses with the marker protein SynapsinI using primary antibodies followed by secondary antibodies conjugated either with DNA docking strands or with Alexa488 dye. The SynapsinI signals from Exchange-Confocal and from conventional IF were obtained with 561 nm and with 488 nm excitation, respectively. We observed colocalization of fluorescence signals from these two methods with a correlation coefficient of 0.96. We also performed Exchange-Confocal based co-localization analysis of SynapsinI and Synaptophysin, both of which are present in synaptic vesicles (**Fig. 2b**). We obtained a correlation coefficient of 0.80, which is similar to values that have been reported using array tomography^20^.

**Figure 2.**
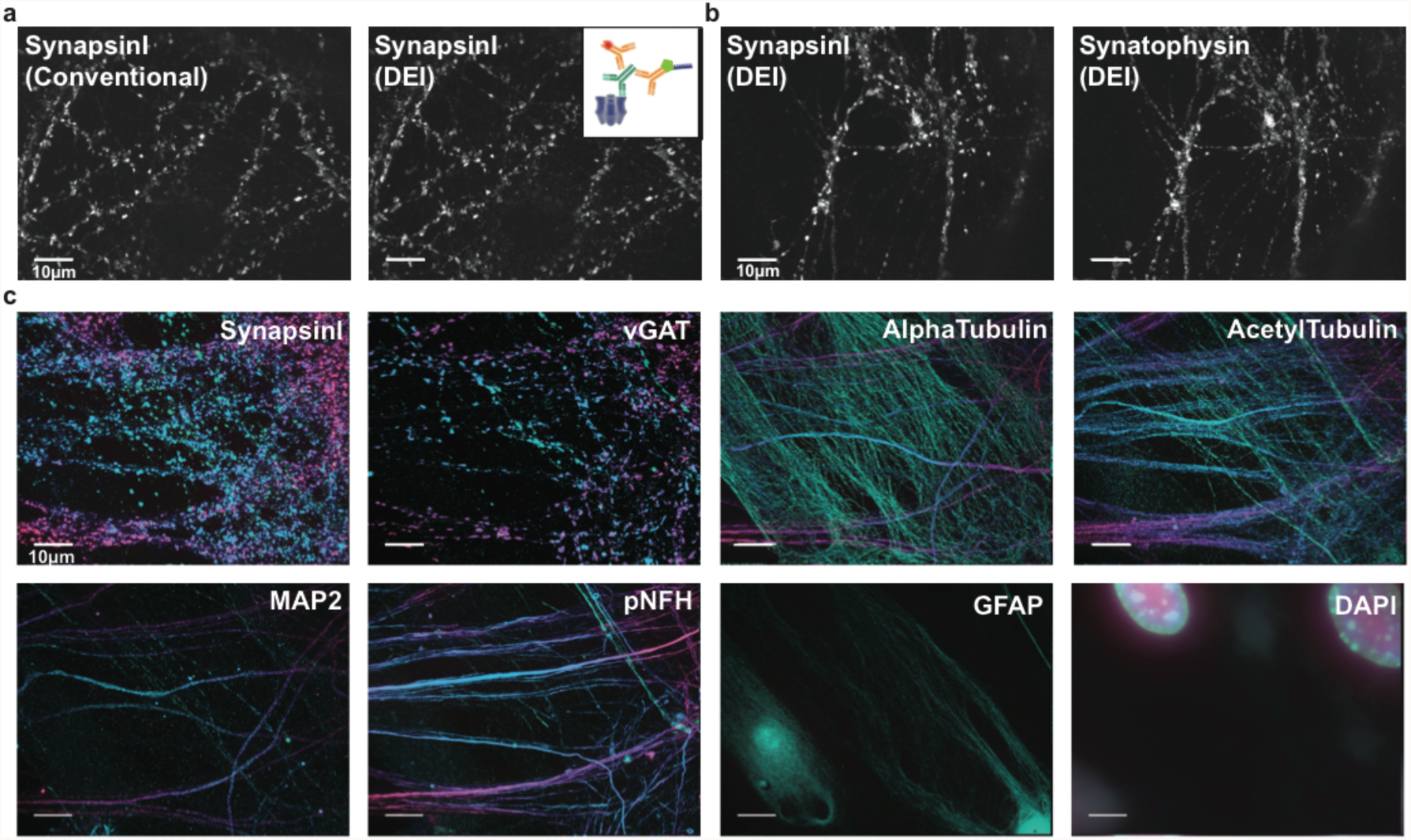
Multiplexed diffraction-limited confocal imaging with DEI. **(a)** Comparison of conventional staining using dye-conjugated antibodies and DEI using DNA-conjugated antibodies. Fixed neurons were stained with primary antibodies targeting SynapsinI, followed by both Alexa647-conjugated and DNA-conjugated antibodies, as shown in the schematic. DNA-conjugated antibody signals were visualized using Cy3b-imager strands. The correlation coefficient of the two images was 0.96. (**b**) Colocalization of SynapsinI and Synaptophysin in neurons visualized using two rounds of DEI. (**c**) Multiplexed eight-target imaging in neurons. Fixed DIV (Days *in vitro)* 14 mouse hippocampal neurons were stained with DNA-conjugated antibodies against SynapsinI, vGAT, MAP2, pNFH, GFAP, AlphaTubulin and AcetylTubulin. A 3D image stack of 14 μm thickness in z-axis was taken for each target and displayed as 2D color-coded maximum intensity projection (bottom to top: green to red). Scale bars: 10 μm. DNA docking strand sequences are listed in **Supplementary Table 4**.

As Exchange-Confocal requires sequential application of imager strands labeled with the same fluorophore, efficient imager strand removal is critical. We tested the change in fluorescence intensity between each cycle of imager strand exchange (**Supplementary Fig. 4**). DIV14 mouse hippocampal neurons were fixed and stained with antibodies against glial fibrillary acidic protein (GFAP, a marker protein for astrocytes) and beta3Tubulin (a marker protein for neurons). P1* and P2* imager strands were sequentially applied to visualize GFAP and beta3Tubulin, respectively. The fluorescence intensity after washing with PBS was decreased to the background level and thus negligible compared to signal levels in the other images, confirming sufficiently efficient removal of imager strands from the solution.

### Multiplexed diffraction-limited Exchange-Confocal imaging in primary neuron cultures

To demonstrate multiplexed Exchange-Confocal, we next imaged eight targets in a fixed primary mouse hippocampal neuron culture (**Fig. 2c and Movie 1**). SynapsinI antibodies were used to mark all synapses, and Vesicular GABA transporter (vGAT) antibodies labeled inhibitory synapses. Five other structural proteins were also labeled, including Microtubule associated protein2 (MAP2) (a dendritic marker), phosphorylated neurofilament heavy chain (pNFH) (in neurites), AlphaTubulin (microtubule component), AcetylTubulin (microtubule component) and GFAP (astrocyte marker). DAPI was used to stain nuclei. For the eight protein targets, we performed dual-color imaging (using Cy3b- and Atto655-conjugated imager strands) to reduce probe exchange cycles. Sample drift was monitored by the signals from the 488 nm channel, and images were registered accordingly (**Supplementary Fig. 5**).

Three-dimensional (3D) images were taken for each target using a spinning disk confocal microscope and the color-coded 2D maximum projection images were displayed for each target (**Fig. 2c**). We used green and red colors to represent the signals from the bottom and top focal planes respectively. A color gradient from green to red was then used to represent the signals from intermediate focal planes. Astrocytes, labeled with GFAP, were mostly shown in green, consistent with the fact that astrocytes grew beneath neurons. SynapsinI labeled both excitatory and inhibitory synapses, while vGAT only labeled inhibitory synapses. As expected, SynapsinI signals were more abundant than those of vGAT. AlphaTubulin were observed in both astrocytes and neurons across the whole z-stack. Interestingly, acetylTubulin signals were more abundant in neurons than in astrocytes.

### Multiplexed diffraction-limited Exchange-Confocal imaging in thick tissue sections

To test the applicability of Exchange-Confocal to tissue samples, we performed eight-target Exchange-Confocal in fresh-frozen mouse retina tissue sections (**Fig. 3a** and **b**). We chose retina samples because the tissue organization has been intensively studied and different cell types can be distinguished using protein markers^21^’^22^. A 40 μm thick retina section was stained using DNA-conjugated antibodies against SV2, GFAP, Cone arrestin, Chx10, Vimentin, and Synapsin, and imaged with six rounds of exchange using Cy3b-conjugated imager strands. Lectin-Alexa488 was used to stain blood vessels and imaged for every exchange cycle for image registration. DAPI was used to stain the nucleus. As expected, every protein species was truthfully detected using Exchange-Confocal with the distribution of each target being consistent with previous reports^21,22,23^. SV2 and Synapsin are both located in synapses. SV2 exists in both Outer Plexiform Layer (OPL) and Inner Plexiform Layer (IPL), whereas Synapsin is only located in the IPL, similar to what has been reported in Salamander retina^24^ (**Fig. 3b**). It should be noted that SynapsinI antibody was used to stain the sample and the lack of Synapsin signal in the OPL only reflects the absence of SynapsionI, which could be replaced by alternative forms of Synapsin, such as Synapsin II or III. GFAP marks astrocytes that are located close to the Ganglion Cell Layer (GCL) and Muller glial endfeet. Cone arrestin marks the cone photoreceptor cells in the Outer Nuclear Layer (ONL). Vimentin labels Muller cells that spread across multiple layers. Chx10 is a pan-bipolar cell marker^23^ located in the Inner Nuclear Layer (INL). Another five-target Exchange-Confocal experiment was performed on a 10 μm thick formaldehyde fixed mouse brain section (**Supplementary Fig. 6**).

**Figure 3.**
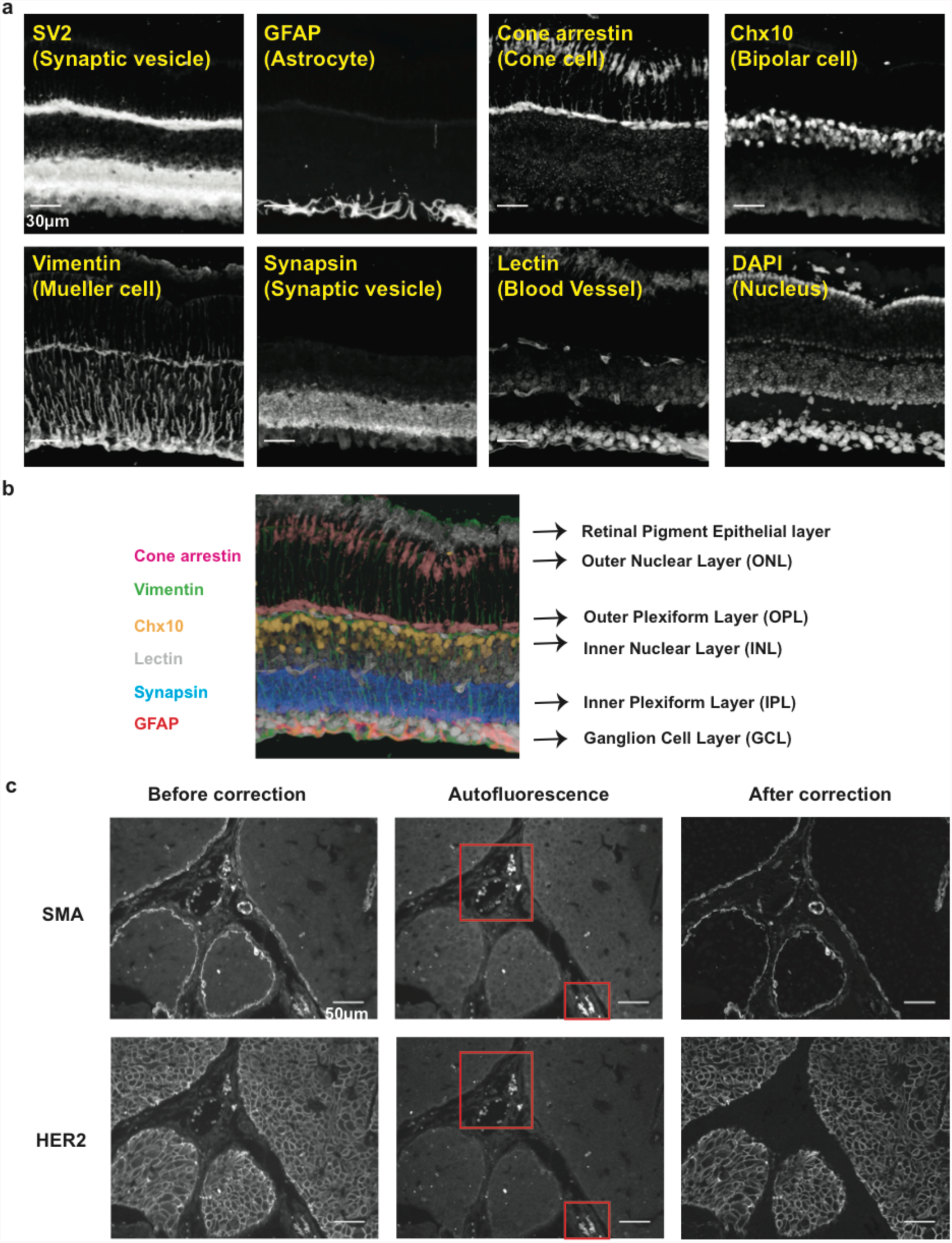
Multiplexed DEI of tissue samples. **(a)** A 40 μm thick fresh frozen mouse retina section was stained with antibodies targeting SV2, GFAP, Cone arrestin, Chx10, Vimentin, Synapsin. 3D images were taken with six rounds of exchange of Cy3b-labeled imager strands. Blood vessels were stained with Alexa488-conjugated lectin probes and imaged in every exchange cycle for image registration. The nucleus was stained with DAPI. Scale bars: 30 μm. (**b**) Merged six-target image reveals different layers of cells in the retina. (**c**) Autofluorescence correction with DEI on a paraffin-embedded breast tumor section. Autofluorescence images were taken before adding imager strands with the same laser intensity and camera exposure time, and then subtracted from the corresponding target images to obtain autofluorescence-corrected images. Note that the strong autofluorescence (presumably from blood cells, labeled with red square) was eliminated in the corrected images. Scale bars: 50 μm. DNA docking strand sequences are listed in **Supplementary Tables 5** and 7.

We also tested Exchange-Confocal in paraffin-embedded tissue samples, and performed two rounds of probe exchange to visualize HER2 and smooth muscle actin (SMA) in a 4 μm formalin-fixed and paraffin-embedded intraductal breast tumor carcinoma tissue from a HER2+ xenograft of SUM225 tumor cells (**Fig. 3c**). SMA stains the myoepithelial cells surrounding the intraductal tumor as well as stromal fibroblasts. We also note that Exchange-Confocal permits simple autofluorescence correction, an additional advantage over conventional fluorescence imaging for tissue samples. Autofluorescence, caused by the presence of various endogenous molecules (e.g. reduced NAD(P)H, flavins, reticulin fibers, lipofuscins, elastin and collagen) can mask true target signals^25^. Although a few approaches have been developed, such as autofluorescence quenching using Sudan Black B, photobleaching with high intensity lasers and post-measurement image correction using complex mathematical models, they require optimization specific for each type of sample and/or may cause sample damage if harsh treatment is performed^25^. When performing DEI – as fluorophore-tagged imager strands are not added until the sample is ready to be imaged on the microscope – an image exhibiting only autofluorescence can be acquired immediately before the addition of imager strands, and subsequently subtracted from the true target image. In **Figure 3c**, autofluorescence signals were captured before addition of imager strands in the same field of view. Compared with images before correction, the ‘false’ signals indicated by the red arrows were significantly reduced in the corrected images. It should be noted that the laser intensity and camera exposure time for autofluorescence images should be identical to those used for the real target image to ensure accurate correction.

### Multiplexed Exchange-SIM and -STED imaging in neurons

Although diffraction-limited Exchange-Confocal enables faster and deeper sample imaging, its resolution may not be sufficient to address certain biological questions that require subcellular resolution. To achieve this, we applied DNA-Exchange-Imaging to various fast SR imaging microscopy platforms. First, we performed DEI using structured illumination microscopy (SIM), which doubles the achievable resolution^26^. Here in Exchange-SIM, we stained BSC1 cells with antibodies against AlphaTubulin followed by DNA-conjugated secondary antibodies (**Fig. 4a and b**). We measured the full width at half maximum (FWHM) of microtubules by Gaussian fitting the intensity plot of 20 microtubule crosssections, and obtained an average of ~2-fold reduction of FWHM, consistent with the theoretical resolution enhancement for commercial SIM microscopes (**Fig. 4c and d** and **Supplementary Table 11)**. While improving spatial resolution helps to resolve fine molecular structures, it also renders the experiment more sensitive to sample drift during buffer exchange process. To reduce drift-induced errors, we adapted a phase correlation-based algorithm^27^ to perform subpixel registration (see Methods for more details). The algorithm correctly identified sample drift between different exchange cycles and registered images accordingly (**Fig. 4e**). Multiplexed SIM imaging was performed by four rounds of exchange with Cy3b-conjugated imager strands targeting alphaTubulin, Vimentin, Tom20, and betaTubulin (**Fig. 4f**). An upsampling factor of 5 in x- and y-axis and a factor of 2 in z-axis was used to perform subpixel image registration, resulting in a subpixel precision of 5 nm in x and y-axis and 75 nm in z-axis.

**Figure 4.**
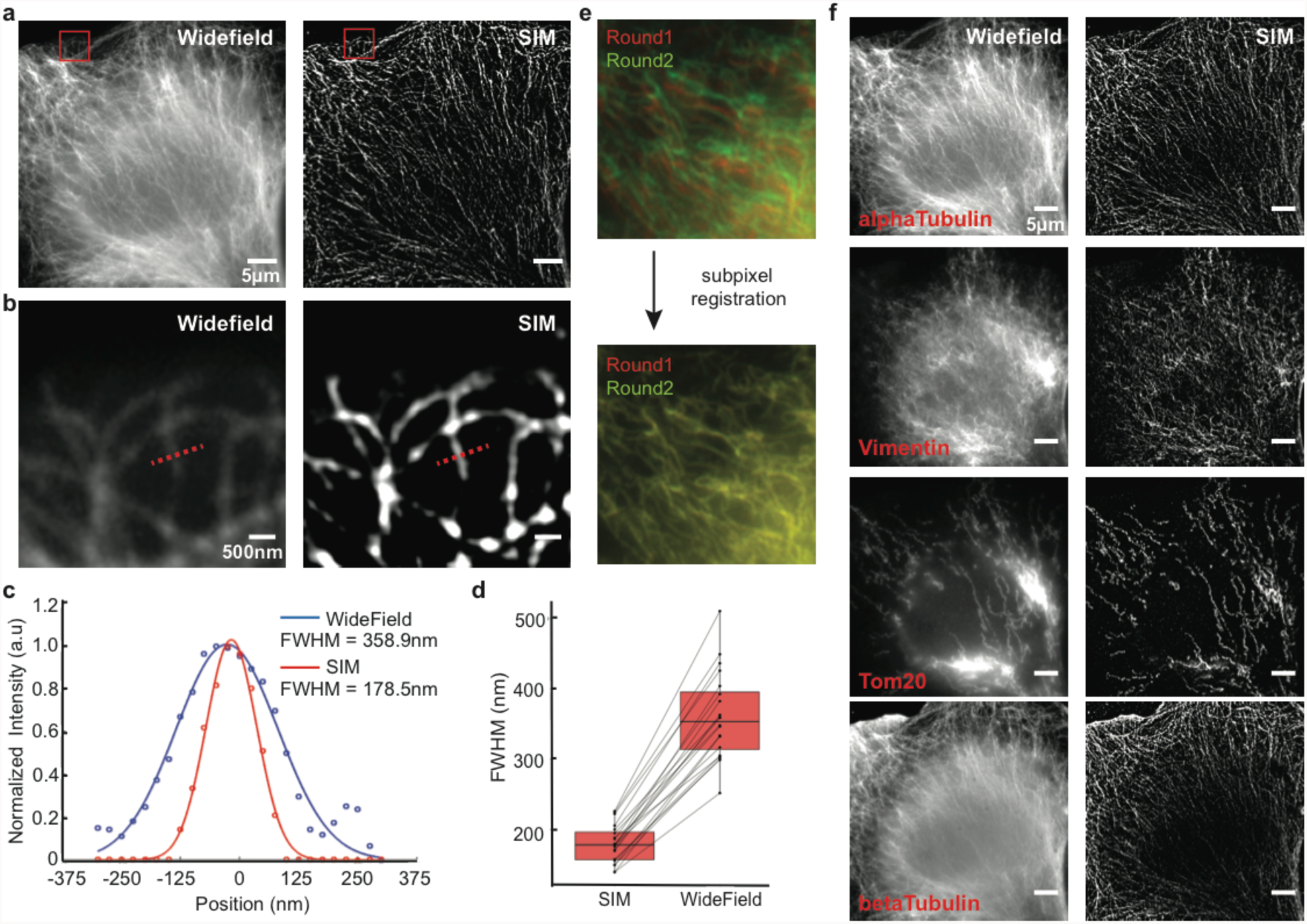
Four-target DEI with SIM in BSC1 cells. **(a)**comparison of widefield and SIM images on alphaTubulin. Scale bars: 5 μm. **(b)** Zoom-in views of the microtubules highlighted by red square in **a. (c)** Measurement of the apparent width of microtubules using Full width at half maximum (FWHM) criterion. The intensity plot of the cross-section highlighted in **b** were fitted using a Gaussian. **(d)** FWHM measurement of 20 microtubule cross-sections revealed 2.014 ± 0.045 fold reduction of FWHM (the error range is SEM; boxes denote median values ± quartiles). **(e)** Subpixel registration of images in different exchange rounds. Vimentin was stained with both DNA-conjugated and Alexa488-labeled antibodies, with the 488 nm channel used for image registration. (f) Multiplexed 3D Exchange-SIM imaging in BSC1 cells. The 2D maximum intensity projections are presented here. Scale bars: 5 μm. DNA docking strand sequences are listed in **Supplementary Table 8.**

A similar multiplexed experiment was done using a STED microscope (**Supplementary Fig. 7**). Together with previous related Exchange-STED work applied to synthetic DNA nanostructures^28^, our results show that DEI is generally compatible with SIM and STED microscopy and can be used for rapid multiplexed SR imaging.

### Multiplexed Exchange-PAINT imaging in neurons

For even higher spatial resolutions, we turned to our previous Exchange-PAINT^17^ method and demonstrated eight-target super-resolution imaging in cultured neurons. DIV14 mouse hippocampal neurons were fixed and stained with antibodies against AcetylTubulin, AlphaTubulin, Vimentin, Tom20, SynapsinI, Bassoon, vGAT, and Gephyrin, utilizing our recently developed DNA-antibody labeling chemistry^19^. While synapsin1 and vGAT antibodies label all and inhibitory synaptic vesicle clusters, respectively, Bassoon is a marker for the presynaptic active zone and gephyrin marks postsynaptic scaffolds at inhibitory synapses. AcetylTubulin and AlphaTubulin are both microtubule components. Vimentin is a component protein in intermediate filaments, and Tom20 is located in the mitochondria. Eight rounds of Exchange-PAINT imaging with Atto655-conjugated imager strands were performed to visualize each target (**Fig. 5**).

**Figure 5.**
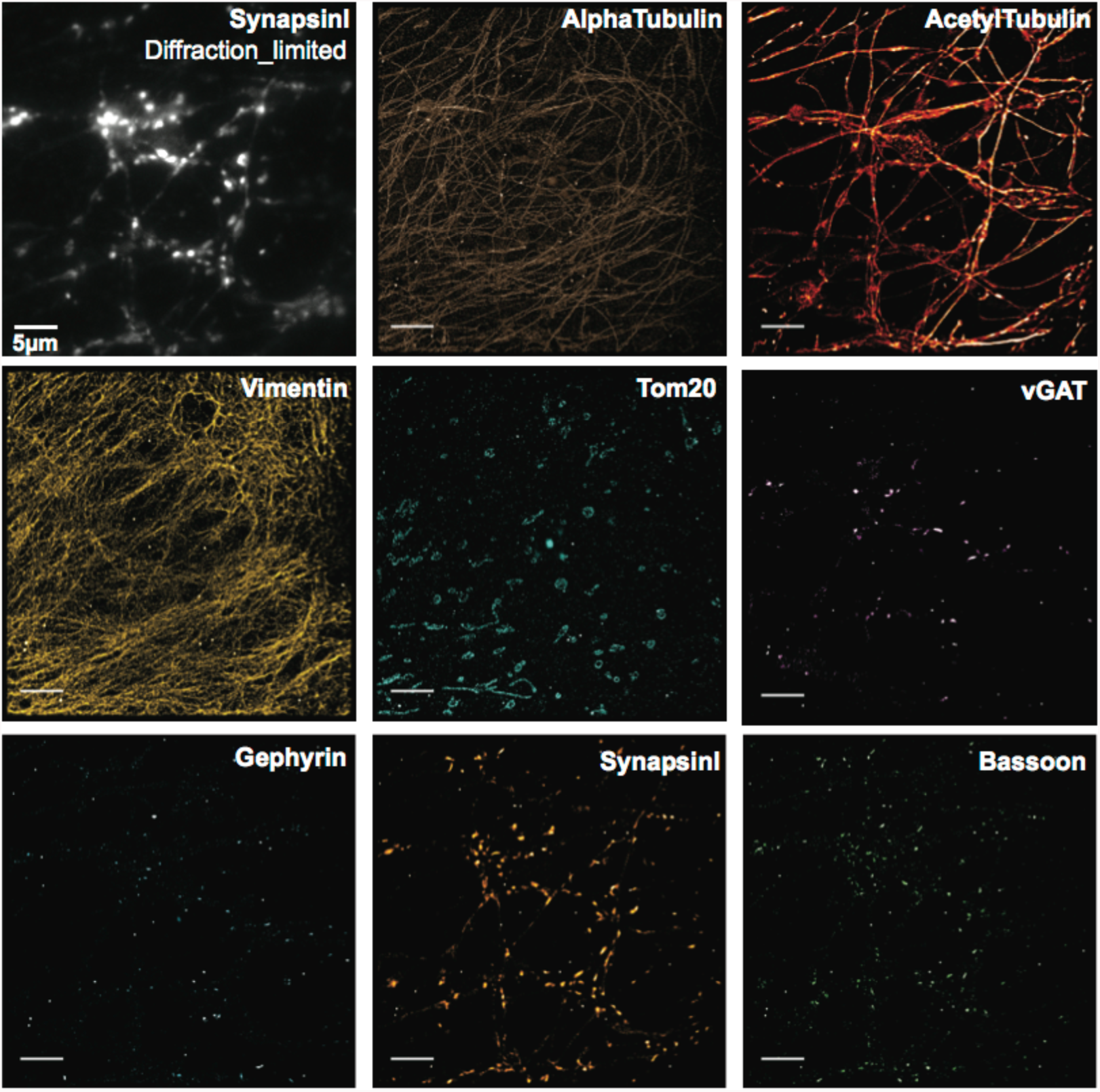
Eight-target chromatic aberration-free Exchange-PAINT imaging in primary neurons. Fixed DIV14 mouse hippocampal neurons were stained with DNA-conjugated antibodies targeting AlphaTubulin, Vimentin, vGAT, Gephyrin, SynapsinI, Bassoon, AcetylTubulin, and Tom20. SynapsinI was additionally labeled with Alex488-conjugated secondary antibodies for selecting regions of interest. In total, eight rounds of Exchange-PAINT imaging with Atto655-conjugated imager strands were performed, visualizing all targets. Scale bar: 5 μm. DNA docking strand sequences are listed in **Supplementary Table 10.**

To demonstrate the improvement in resolution, we compared the quality of diffraction-limited and SR images for microtubules, and merged Bassoon and Gephyrin from **Figure 5** (**Supplementary Fig. 8** and **9**). Individual microtubule filaments were clearly resolved in the SR image but not in the diffraction-limited image (**Supplementary Fig. 8a** and **b**). A region was selected for a magnified view with two microtubules in close proximity to each other, and the distance between the two filaments was measured to be 108 nm **(Supplementary Fig. 8c** and **d**). In the Bassoon and Gephyrin merged image, presynaptic Bassoon signals can be distinguished from the postsynaptic Gephyrin signals in the SR image but not in the diffraction-limited image (**Supplementary Fig. 9**).

One key application of multiplexed imaging is to detect protein-protein tight co-localization. To test the applicability of Exchange-PAINT for such studies, we merged the four synaptic protein images from **Figure 5** to assay co-localization of these proteins (**Fig. 6a**). We first compared the diffraction-limited and super-resolution images. Individual synapses are difficult to distinguish from each other in the diffraction-limited images but can be clearly visualized in the super-resolution images (**Fig. 6b**). Particularly, the synapse orientation can be detected by lining synapsin (synaptic vesicle marker that is further from the presynaptic membrane), Bassoon (active zone marker that is closer to the presynaptic membrane) and gephryin (postsynaptic density marker on the postsynaptic sites) (**Fig. 6b**). We also selected one region for a magnified view (**Fig. 6c**). SynapsinI and Bassoon are known to be present in both excitatory synapses and inhibitory synapses, whereas vGAT and Gephyrin selectively label inhibitory synapses^20^. Three synapses were included in this region. Two of them contained only SynapsinI and Bassoon signals, suggesting they were excitatory synapses, whereas the middle synapse contained all four targets, indicating that it was an inhibitory synapse (**Fig. 6c**). SynapsinI, Bassoon and vGAT were present in the presynaptic site and therefore well separated from the signal from Gephyrin that existed in the postsynaptic site. The distribution patterns of SynapsinI and vGAT, both of which were localized on synaptic vesicles, correlated well with each other. The result indicates Exchange-PAINT is well suited for high-resolution visualization of protein-protein tight co-localization *in situ*.

**Figure 6.**
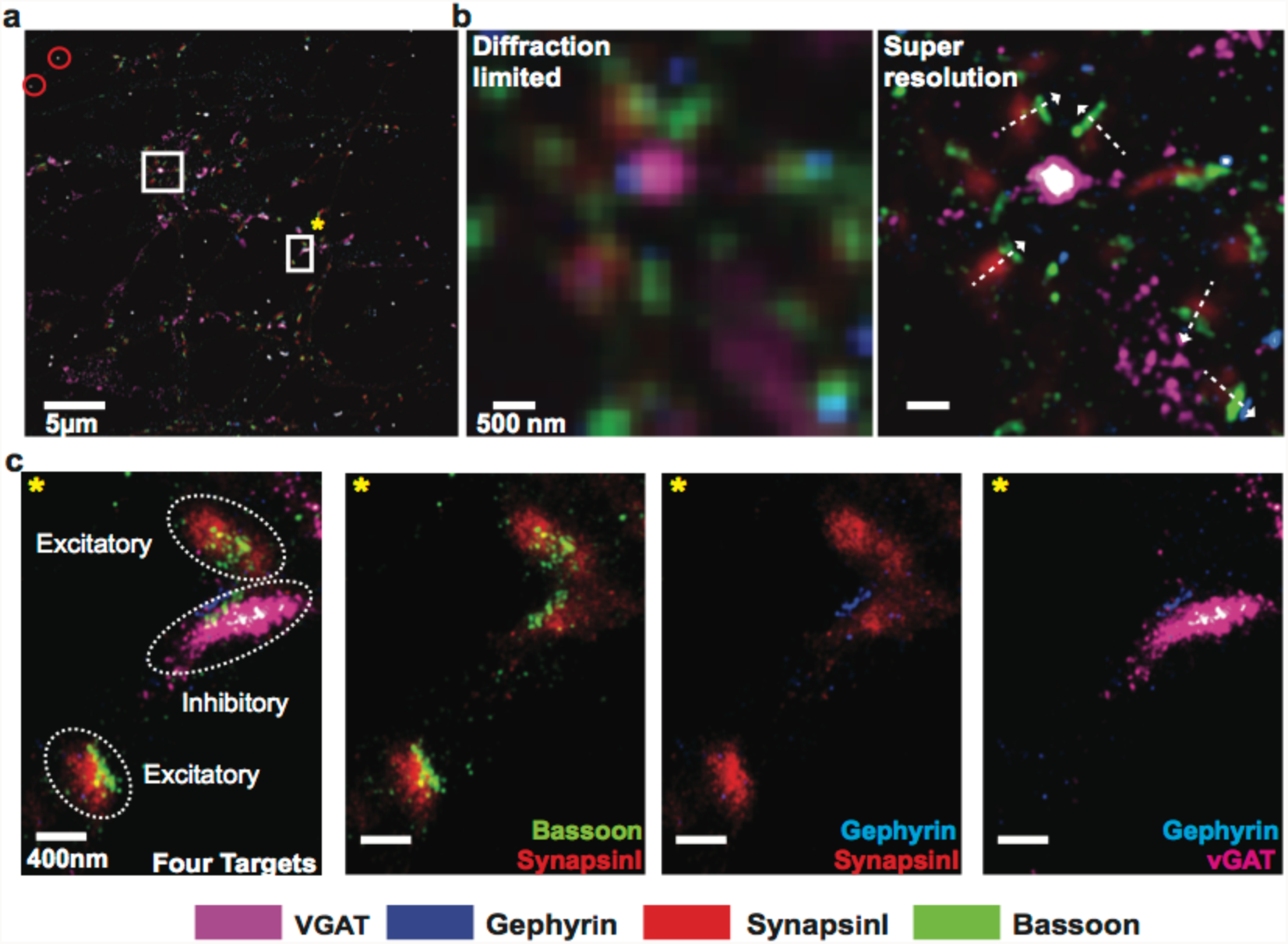
Co-localization of synaptic proteins using multiplexed Exchange-PAINT imaging. **(a)** The images of synaptic proteins from **Figure 5** were merged using gold nanoparticles as registration markers (highlighted with red circles). Scale bar: 5 μm. **(b)** Comparison of diffraction-limited and super-resolution images of four synaptic proteins from the region highlighted with a white square without *. The orientation of synapses could be visualized in super-resolved image as indicated by the white dashed arrows. Scale bar: 500 nm. **(c)** One region from **a** was selected for a magnified view (highlighted with a white square with *). Scale bars: 400 nm.

## Discussion

An increasing body of effort has been devoted to molecular heterogeneity mapping in single cells. Such *in situ* ‘omics’ studies, including transcriptomics and proteomics, have the potential to greatly expand our knowledge about how cells and tissues are organized to realize their biological functions. Several techniques, such as laser capture microdissection-assisted single-cell RNA sequencing^29^, Fluorescence in situ RNA sequencing^30^, highly multiplexed FISH^31^, have been developed to enable spatially-resolved transcriptomics. *In situ* proteomics analysis, on the other hand, has not been widely performed mainly due to the lack of efficient and practical methods, particularly as existing sequential IF imaging methods require multi-rounds of time-intensive immunostaining. DEI herein provides a simple, efficient, and versatile tool to map diverse proteins *in situ* with flexible choice regarding achievable spatial resolution. It has multiple advantages: (1) DEI allows fast multiplexed data acquisition and probe exchange, as targets are simultaneously immunostained and transient binding of imagers to docking strands is rapid (minutes); (2) DEI – as other sequential imaging approaches – allows image acquisition with a single laser line, thus avoiding chromatic aberration (**Supplementary Fig. 10**) and eliminating time-consuming optimization of imaging setting (*e.g.* immersion medium for SIM) for individual laser channels; (3) DEI allows straightforward re-imaging of earlier targets. This allows users to rapidly scan a sample *e.g.* using fast Exchange-Confocal to determine regions of interest, and then re-probe these with shorter imager strands for higher resolution imaging *e.g.* using Exchange-PAINT; (4) DEI does not require specialized instruments (e.g. mass spectrometers for MIBI) or harsh buffer treatment (*e.g*. acidified KMnO_4_ or H_2_O_2_) to quench fluorescence signals. The labeling protocols and imaging instruments are identical to standard and well-established immunostaining methods, the only difference being the use of a DNA-tagged antibody as opposed to a dye-tagged antibody, thus making it easily accessible to common biological labs.

A key requirement for sequential imaging is to minimize sample drift during an experiment. All of our buffer exchange experiments were performed without removing samples from the microscope stage. A fluidic chamber system has been described in our original Exchange-PAINT paper^17^ and can be used to reduce the physical disturbance caused by buffer exchange. A registration marker channel, either bright field or other fluorescence channels, is required to record sample drift for post-experiment image registration. In this current study, we adapted a subpixel registration algorithm^27^ that can perform translation drift correction with a user-defined up-sampling factor. It increased the registration accuracy, which is important when super-resolution imaging is performed. Z-axis drift can be easily managed by using commercially available focus maintaining systems.

We note that we occasionally observe non-specific nuclear staining from DNA-conjugated primary antibodies, which is likely an antibody-specific phenomenon. Interestingly, we did not observe a similar phenomenon for DNA-conjugated secondary antibodies (**Supplementary Fig. 11**). It has been suggested that addition of dextran sulfate to the incubation buffer can alleviate the non-specific binding^32^. Addition of Herring sperm DNA and polyT DNA has also been used to block nonspecific interaction caused by DNA^14^. We also notice that using saponin instead of Triton or Tween as detergent during staining does not permeabilize the nucleus membrane and hence prevents antibodies from entering the nucleus.

In summary, we have developed DNA-Exchange-Imaging as a rapid and versatile multiplexed imaging technique for both diffraction-limited and super-resolution *in situ* imaging in cells and in tissues. The intrinsic programmability of imager/docking strand interaction renders DEI easily adaptable to diverse imaging platforms, including standard resolution Exchange-Confocal demonstrated here, and various super-resolution methods including Exchange-SIM demonstrated here, Exchange-STED demonstrated here and in related work^28^, Exchange-STORM demonstrated in our recent related work using stably attached imager strands^33^, and Exchange-PAINT^17^ demonstrated in our original work with increasing resolution. Beyond these validated imaging platforms, we also expect that DEI is in principle compatible with many other imaging methods. For example, a combination of DEI with ultrathin sectioning of samples could allow correlative light and electron microscopy imaging. Additionally, DEI is also in principle compatible with DNA mediated *in situ* signal amplification methods (e.g. hybridization chain reaction^34^, and rolling circle amplification^35^), potentially permitting rapid, spectrally unlimited multiplexing for low abundance targets. Combination of DEI with tissue clearing methods, such as CLARITY^36^ and SWITCH^37^, would allow imaging of thick tissue sample. Combination of DEI with expansion microscopy^32^ would further allow imaging thick samples with nanoscopic resolution. Integration of DEI with neuron tracing techniques, such as Brainbow^38^, could allow simultaneous detection of neuronal connectivity and underlying molecular characteristics, such as cell identity. The resultant ‘Molecular Connectome’ would complement the ‘Anatomical Connectome’^39^ and help us understand the brain function across multiple scales from circuits to molecules.

## Acknowledgements

We thank X. Chen, F. Schueder, and C. Cepko for helpful discussions. We thank Z. Liang for kindly providing experimental materials. We thank L. Ding from Enhanced Neuroimaging Core in Harvard for assistance in STED microscopy. This work is supported by NIH grants (1U01MH106011, 1R01EB018659), NSF grant (CCF-1317291), and ONR grants (N00014-13-1-0593 and N00014-14-1-0610) to P.Y.; Emmy Noether Fellowship (DFG JU 2957/1-1), ERC Starting Grant (MolMap, Grant agreement number 680241) and support from Max Planck Society to R.J.; Wellcome Trust-DBT India Alliance Intermediate Fellowship (IA/I/16/1/502368) to S.S.A.; NIH grant (RM1HG008525) to G.M.C.; NIH grant (R01NS083898) to P.S.K.; NIH grant (P01 CA080111.) to J.S.B.; NIH grants (1R24MH106075, NIH 1R01NS087950, and NIH 1R01MH103910), and support from HHMI-Simons Faculty Scholars Program, the Open Philanthropy project, the MIT Media Lab, and the New York Stem Cell Foundation to E.S.B. Y.W. acknowledges support from the Chinese Scholarship Council. M.S.A. and M.D. acknowledge support from HHMI International Student Research Fellowships. S.S.H.W. acknowledges support from NSF fellowship DGE1144152. P.W.T. acknowledges support from the Hertz Foundation.

